# Mining underutilized whole-genome sequencing projects to improve 16S rRNA databases

**DOI:** 10.1101/2021.01.01.425045

**Authors:** Ben Nolan, Florence Abram, Fiona Brennan, Ashleigh Holmes, Vincent O’Flaherty, Leighton Pritchard, Nicholas R. Waters

## Abstract

Current approaches to interpreting 16S rDNA amplicon data are hampered by several factors. Among these are database inaccuracy or incompleteness, sequencing error, and biased DNA/RNA extraction. Existing 16S rRNA databases source the majority of sequences from deposited amplicon sequences, draft genomes, and complete genomes. Most of the draft genomes available are assembled from short reads. However, repeated ribosomal regions are notoriously difficult to assemble well from short reads, and as a consequence the short-read-assembled 16S rDNA region may be an amalgamation of different loci within the genome. This complicates high-resolution community analysis, as a draft genome’s 16S rDNA sequence may be a chimera of multiple loci; in such cases, the draft-derived sequences in a database may not represent a 16S rRNA sequence as it occurs in biology. We present Focus16, a pipeline for improving 16S rRNA databases by mining NCBI’s Sequence Read Archive for whole-genome sequencing runs that could be reassembled to yield additional 16S rRNA sequences. Using riboSeed (a genome assembly tool for correcting rDNA misassembly), Focus16 provides a way to augment 16S rRNA databases with high-quality re-assembled sequences. In this study, we augmented the widely-used SILVA 16S rRNA database with the novel sequences disclosed by Focus16 and re-processed amplicon sequences from several benchmarking datasets with DADA2. Using this augmented SILVA database increased the number of amplicon sequence variants that could be assigned taxonomic annotations. Further, fine-scale classification was improved by revealing ambiguities. We observed, for example, that amplicon sequence variants (ASVs) may be assigned to a specific genus where Focus16-correction would indicate that the ASV is represented in two or more genera. Thus, we demonstrate that improvements can be made to taxonomic classification by incorporating these carefully re-assembled 16S rRNA sequences, and we invite the community to expand our work to augment existing 16S rRNA reference databases such as SILVA, GreenGenes, and RDP.

## Introduction

The use of genetic markers for microbial classification has seen explosive growth over the past decade (Liu et al. 2012; Boers, Jansen, and Hays 2019). The 16S rRNA gene is the standard utilised in the assessment of prokaryotic community composition by amplicon sequencing (Fukuda et al. 2016). 16S rRNA has been used for community analysis in diverse environments such as the gut microbiota of cattle and pigs (Avila-Jaime, Kawas, and Garcia-Mazcorro 2018), soil (Santamaria, Parrado, and López 2018), marine environments (Dang and Lovell 2000), and the human gut (Jovel et al. 2016). The success of this method hinges on the presence of the 16S rRNA gene in all domains and its relatively slow rate of base substitution; thus the rDNA regions can be targeted with primers but the amplicon sequences exhibit enough diversity that organisms can be differentiated at the genus or species level (Woese and Fox 1977; Woo et al. 2008).

Microbial genomes have a range of 16S rRNA gene copy numbers (GCNs), from the many Mycobacteria with a single copy to *Photobacterium damselae* Phdp Wu-121 with 21 copies (Větrovský and Baldrian 2013; Stoddard et al. 2015; Acinas et al. 2004). There may be variability between each 16S rDNA copy within an organism (Sun et al. 2013); this can negatively impact 16S rRNA classification in two ways. First, in taxa with low variability, diversity estimates can be skewed by overestimating taxa with higher GCN and underestimating those with low GCN. A trivial example would be a community of two organisms – one with five rDNA copies, one with a single copy; an even sequencing of the community would show a one-to-five abundance ratio. Second, some organisms have sufficient sequence variability between copies that they may be assigned different taxonomic classifications; indeed, certain extremophiles have been reported to possess very high 16S rRNA copy heterogeneity, up to 9.3% sequence variation in some species (Sun et al. 2013); this is well beyond the 97% or 99% clustering thresholds commonly used for community analysis, and clearly beyond the zero-radius OTU boundary (Edgar 2018).

These intergenomic rDNA repeats complicate community analysis, but each instance of the 16S rRNA contains valuable information. An ideal community analysis framework would utilize a database incorporating this information to both correct for copy number variation between organisms in a community, and correctly relate 16S rRNA variants to each organism.

Amongst the most widely used 16S rRNA databases for bacteria and archaea are Greengenes (DeSantis et al. 2006), SILVA (Quast et al. 2012), and the Ribosomal Database Project (RDP) (Cole et al. 2005). Each contains 16S rRNA sequences derived from multiple major international nucleotide sequence databases, principally EMBL/ DDBJ and Genbank. The databases differ in their approach to sequence classification. The RDP database uses the RDP classifier to assign taxonomy to 16S rRNA sequences (Wang et al. 2007). SILVA and Greengenes inherit a sequence’s taxonomic assignment from the source database (such as NCBI or EBI). SILVA provides a non-redundant database version in which no taxonomic classification contains sequences with greater than 99% pairwise identity (Quast et al. 2012). Although each database performs sequence quality checks, only Greengenes actively checks for chimeric sequence, which can negatively affect 16S taxonomic assignment (DeSantis et al. 2006).

The National Centre for Biotechnology Information (NCBI) provides multiple databases, including the Sequence Read Archive (SRA) (Kodama, Shumway, and Leinonen 2011) for raw high-throughput sequencing data, and the Genome database as an umbrella for draft and complete genomes. Not all genome sequences in the NCBI Genome database have publicly available raw data in the SRA, and only 10% of genomes in the database are closed or complete (Waters et al. 2018). Tabulating the accession types of the SILVA 132 database shows that 9.5% of sequences come from draft genome assemblies; the vast majority (87%) are obtained as amplicon sequences (usually Sanger sequenced), and the remaining 2% come from complete genomes. While Sanger-sequenced amplicons are generally very accurate, a common weakness of draft assemblies from short-read sequencing is incorrect assembly of repeated rDNA regions of a genome, which may be collapsed/merged into a single rDNA. The resulting 16S rRNA sequence could in turn be incorporated into SILVA, GreenGenes, or RDP. This compromises the quality of 16S rRNA databases, and such sequences should be treated with caution. Genome assemblies from short reads are prone to errors in rDNA regions, as the length of the repeated region exceeds read lengths. PCR spanning the rDNA region, followed by Sanger sequencing, or the use of long-read technologies such as PacBio or Nanopore sequencing, can resolve these multiple copies but, as the majority of the data generated over the last two decades comes from short read sequences, fixing collapsed regions remains a valuable goal (Land et al. 2015; Wagner et al. 2016).

The correct re-assembly of multiple rDNA regions of draft genomes can be achieved using riboSeed, which uses a reference genome to help assemble the rDNA regions of a draft genome (Waters et al. 2018). riboSeed exploits the observation that the flanking regions of the rDNA region are highly conserved within a taxon yet variable between rDNA copies in the same genome, by using targeted subassembly to correctly place each re-assembled copy of multi-locus rDNA repeats. The knock-on effect of assembling multi-copy rDNA operons is acquiring highly-accurate 16S rRNA sequences, which can be incorporated into 16S rRNA databases.

Here, we present the results of using such an approach to augment existing 16S rRNA databases with newly-assembled sequences from the SRA data corresponding to pre-existing draft genomes. The additional sequences provide greater coverage of ASVs in publicly-available datasets, aiding efforts to understand microbial communities.

## Methods

### The Focus16 Pipeline

We developed Focus16: a pipeline to augment existing 16S rRNA databases by mining the SRA database for candidate whole-genome sequencing studies for re-assembly. Candidate SRAs are identified, downloaded, subjected to automated quality control, re-assembled with riboSeed to resolve the rDNA operons, given a taxonomic assignment with Kraken, and formatted for addition to existing databases. Kraken2 assigns taxonomy using exact matches to a lowest common ancestor for k-mers from the whole genome assembly, mitigating the risk of misclassification compared to using a 16S rRNA classifier alone.

The pipeline is shown as a flowchart (Figure 1). Details of third-party tools used in Focus16 can be found in the Supplementary Methods section “Third-party software.”

**Figure 1:**
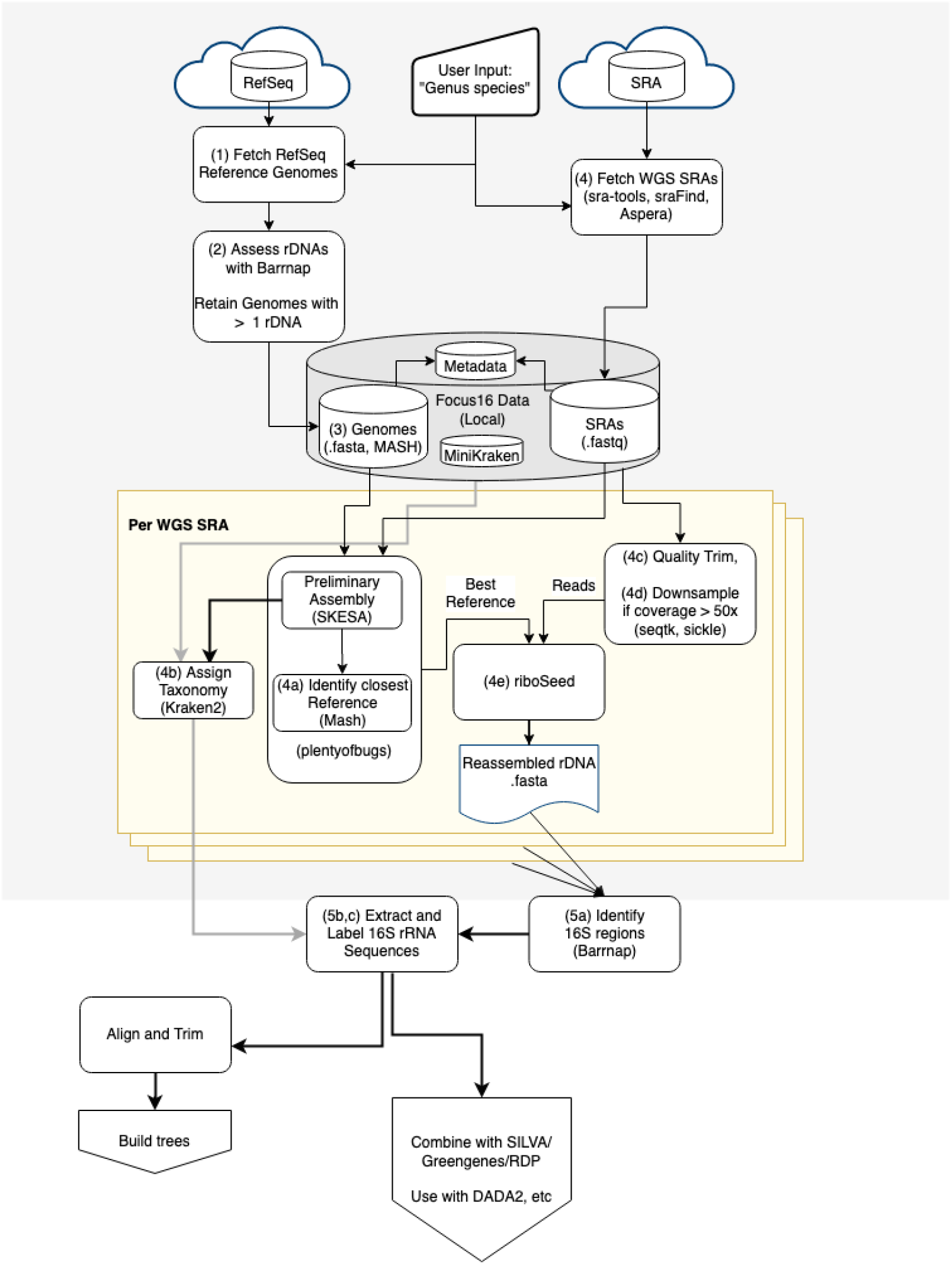
Flowchart of the pipeline that resolves multi-copy 16S loci from sequenced genomes with reads in SRA (as implemented in Focus16). Candidate reference genomes are downloaded from RefSeq. Reads for each SRA are downloaded and Kraken2 is used to assign taxonomy. Corresponding reference genomes and SRA read sets are identified (using SKESA and Mash), and a new assembly constructed from these using riboSeed to resolve 16S rDNAs. The assembled 16S rRNA regions are then taken forward for phylogenetic reconstruction, or to supplement existing reference databases. Numbers refer to stages outlined in the text; gray lines signify to taxonomic information, and black lines signify to sequence information.

Given a genus or “Genus species” binomial, the pipeline progresses as follows:

1. Candidate reference genomes are either provided or are identified and downloaded from RefSeq by matching the provided organism name.
2. Barrnap (Seemann 2020) is used to screen these complete genomes by estimating the 16S rRNA count. Reference genomes with a single 16S rRNA are discarded. This catches two cases:

a. An organism may only have a single 16S rRNA. In this case riboSeed assembly will not improve on existing draft genomes.
b. A draft genome may have been incorrectly attributed the classification of “complete,” and present as having a single rRNA sequence when it in fact has more than one. Any such errant reference genomes are therefore discarded, and the remaining references are available for use in the pipeline.
3. A Mash (Ondov et al. 2016) sketch is generated from the references passing the filtering in step 2.
4. sraFind (https://github.com/nickp60/sraFind) is used to identify all whole-genome sequencing SRA accessions for the organism of interest; these are downloaded with fastq-dump or fasterq-dump (“Sra-Tools” 2019). Steps 4a-4e are applied to each SRA.

a. **Identify closest reference genome**. For a given SRA, the most compatible reference genome is determined via plentyofbugs (Waters 2019), which performs an initial assembly with the fast and highly-accurate assembler SKESA (Souvorov, Agarwala, and Lipman 2018) using a subsample of 1M reads. Mash is used to identify the closest match between the preliminary assembly and all the reference genomes from step 1. If no close match above a user-defined threshold (defaulting to a Mash distance of 0.1, roughly corresponding to a within-genus match (Ondov et al. 2016)) is identified, the SRA is skipped; otherwise, the closest match is retained for use as a reference genome.
b. **Classify Assembly**. Kraken2 (Wood, Lu, and Langmead 2019) is used to assign taxonomic classification to the preliminary SKESA assembly. The highest-ranked binomial name is recorded, and the full Kraken2 report is stored in the output folder for inspection.
c. **Pre-Assembly Quality Control**. Reads are run through several quality control steps. The average length of the reads is checked; an SRA that contains reads of very low length (i.e. an average less than 65bp) is rejected, as very short reads cannot be used effectively by riboSeed to differentiate rDNA flanking regions. Reads are then quality trimmed with Sickle (Joshi and Fass 2011) using default parameters. fastp (Chen et al. 2018) is used to identify and remove any remaining adapter sequences. For paired-end runs, unpaired reads are rejected.
d. **Downsample**. Read coverage is assessed using either a user-provided estimate of genome length or the length of the reference genome. If read coverage exceeds a user-specified threshold (defaults to 50x coverage, as further coverage can artificially support sequencing errors; see (Bankevich et al. 2012)), trimmed reads are down-sampled to reach the desired coverage with seqtk (Li 2020).
e. **De fere novo Assembly**. The SRA reads (or downsampled reads from 4d) are then assembled using riboSeed, using the reference genome determined in (2a) as a template genome. Subassemblies are performed with SPAdes (Bankevich et al. 2012); the default parameters of 3 rounds of seeding and 1kbp flanking regions are used.
5. 16S rDNA sequences are extracted and formatted.

a. Barrnap is run on either the subassemblies (“fast” mode) or the final assembly (“full” mode).
b. Taxonomy assigned by Kraken2 in step 4b is used to label extracted sequences.
c. Sequences are written to a fasta file that matches the format used by the SILVA database.

In our analysis, we filtered to retain only full-length sequences by removing any 16S rRNA under 1358bp (under the 1st quartile of the sequence lengths in SILVA). Additionally, we removed any 16S rRNA for which Kraken2’s report showed inconclusive domain-level taxonomic assignment: assemblies were excluded if a single domain was not assigned to over 70% of contigs. This was done to remove potentially-contaminated datasets (see Supplementary Methods section “Identifying poor taxonomic assignments,” Figures S4 and S5).

### Implementation

The Focus16 pipeline can be installed from PyPI or from the source hosted on GitHub at https://github.com/FEMLab/focus16; all of the dependencies can be easily managed with Conda for reproducibility. The package was designed to efficiently handle the downloading and re-assembly of large amounts of short-read data. Users can use SRA-tools’s prefetch command for faster downloads of SRA data; the re-assembly status of each SRA is recorded in an SQLite database. For those with access to a computing cluster running Open Grid Scheduler, the time-consuming assembly steps can be distributed as array jobs as needed.

The first time the pipeline is used, an automated setup procedure is run to download the required databases for Kraken2 and sraFind.

Throughout, diagnostic information is recorded; if an aspect of the pipeline fails, rerunning the same command will reuse available intermediate results wherever appropriate.

### Selecting suitable test datasets and identifying genera

Three mock communities described in the DADA2 manuscript (Callahan et al. 2016) were selected to assess the utility of Focus16. These communities, named “Extremes,” “HMP,” and “Balanced,” and comprising 27, 21, and 57 members respectively (Schirmer et al. 2015; Kozich et al. 2013; Callahan et al. 2016), were sequenced on an Illumina MiSeq yielding over 500,000 250bp paired-end reads each.

To provide an assessment of real-world usage of Focus16, we used the data generated in the EndoBiota study (Ata et al. 2019)(PRJEB26800): a survey of microbiomes across three body sites of women with and without endometriosis. These datasets are summarized in Table 1.

**Table 1:**
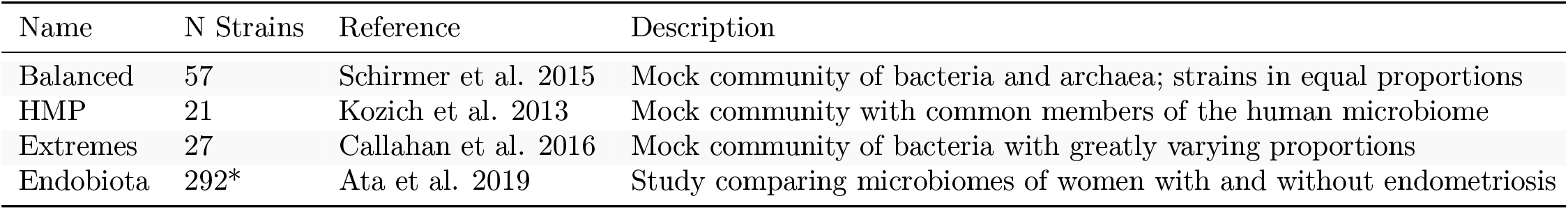
Description of the four datasets considered in this study. The number of strains was readily available for the mock communities; for the Endobiota study, this was determined by a preliminary analysis using DADA2 and SILVA. Asterisk (*) indicates that this is a calculated number, not a value known beforehand.

| Name | N Strains | Reference | Description |
| --- | --- | --- | --- |
| Balanced | 57 | Schirmer et al. 2015 | Mock community of bacteria and archaea; strains in equal proportions |
| HMP | 21 | Kozich et al. 2013 | Mock community with common members of the human microbiome |
| Extremes | 27 | Callahan et al. 2016 | Mock community of bacteria with greatly varying proportions |
| Endobiota | 292* | Ata et al. 2019 | Study comparing microbiomes of women with and without endometriosis |

Unlike the mock communities, the number of genera present in the Endobiota samples is not known *a priori*. We estimated the abundances of community members by processing the samples through DADA2 in a similar manner to how the Balanced, Extremes, and HMP datasets had been analyzed. The analysis script `DADA2_analysis.Rmd` can be found in the supplementary materials repository. Processed data was deposited along with the rest of results generated in this work at https://zenodo.org/record/1172783. In short, DADA2 was used to build error profiles for each of the samples in the study. Reads were then trimmed 30bp on the 5’ end and 40bp on the 3’ end, quality trimmed after two low-Q bases, and any residual phi-X sequence was removed. Merged amplicons were filtered to retain those between 360 and 450 bases. 4.2% of sequences were determined to be chimeric and removed. Taxonomy was assigned to the remaining 16S rRNA sequences using DADA2’s `assignTaxonomy` command with the SILVA non-redundant version 132, and species-level taxonomy was assigned using DADA2’s `addSpecies` command as described in their manual.

In total, 333 unique genera were identified across the four datasets; these were cross-referenced with sraFind and RefSeq, filtering to retain only those with both short-read SRAs available and at least one reference genome for that genus (Supplementary Figure S2, and Supplementary Table S2). The resulting list of 85 genera was used as the input for the pipeline.

### Assembly mode parameter choice

Candidate 16S rRNA sequences for a given organism could be extracted from riboSeed’s subassemblies or the final *de fere novo* assembly, we sought to determine which was the best choice for generating sequence to extend the reference databases. Generating the subassembled sequences alone is a less computationally-intensive process than whole-genome assembly, but the final whole-genome assembly step acts as a further refinement of the subassemblies. To find which source is the best for augmenting 16S rRNA databases, we determined the error (SNP/indel) rates by comparing complete genomes to the sequences recovered from either *de novo* re-assembly of the complete genome, Focus16’s “fast” mode (sequences from riboSeed’s subassemblies only), and “full” mode (sequences from riboSeed’s final *de fere novo* assembly). The *de novo* assemblies were accurate but failed to recover many individual 16S rDNAs, as is expected due to the repeated nature of these regions. riboSeed’s subassemblies have low error rates and successfully reconstruct the most 16S rDNAs (Figure 2), and as such are the ones we report below and recommend for augmenting a database.

**Figure 2:**
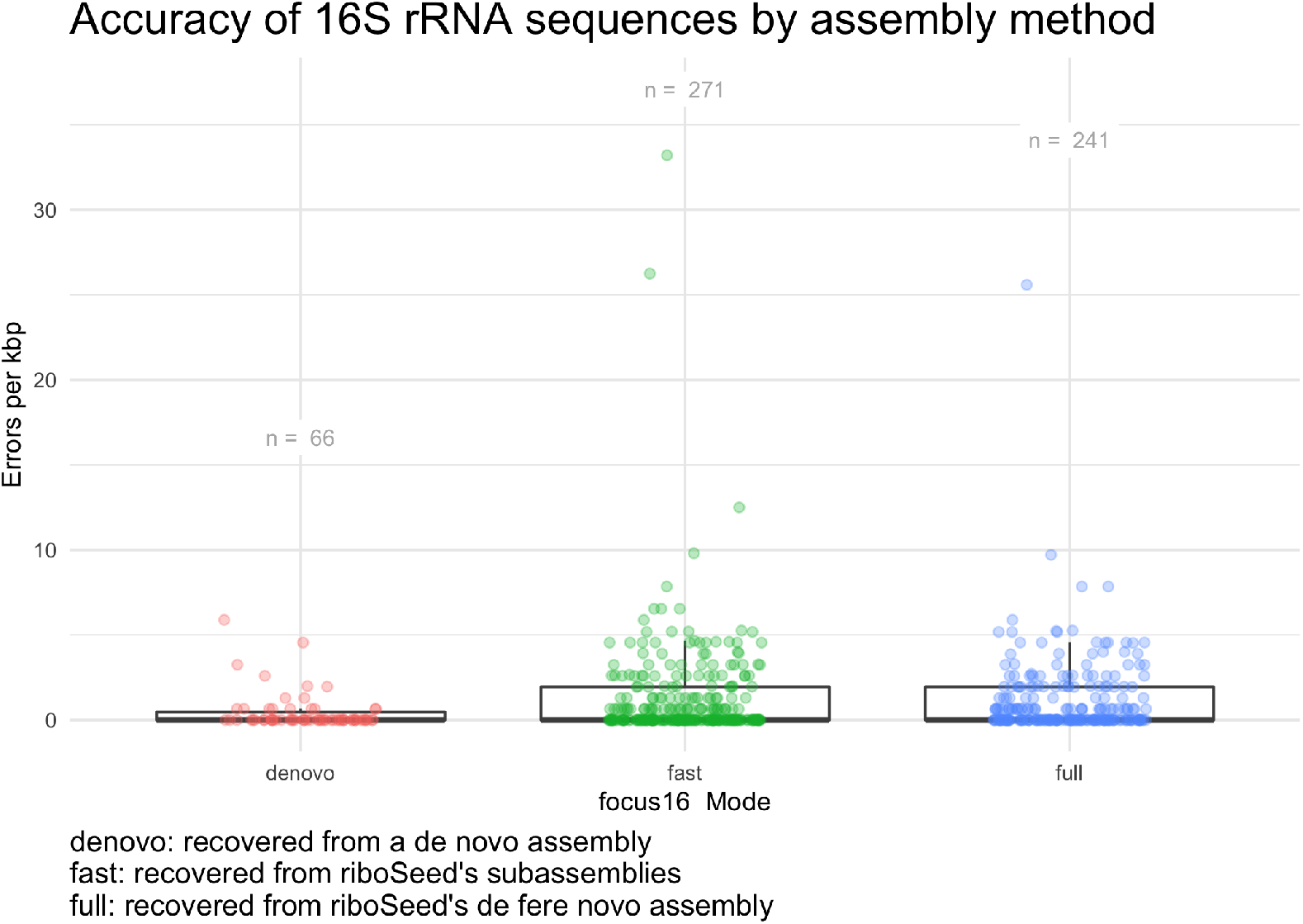
Comparing assembly modes for accuracy. SRAs in our dataset that underwent genome completion were used to identify the most accurate method of 16S rRNA sequence assembly. De novo assembly resulted in highly accurate 16S rRNA sequences, but was only able to recover 66 sequences. ‘–fast’ mode proved to be the best tradeoff between accuracy and efficiency.

### Running Focus16

Focus16 was run in a conda environment (see supplementary repository file “Focus16_env.txt”). Due to computational limitations (namely storage available on the high-performance computing cluster as well as the RAM required for genome assembly), we limited the scope of the analysis to a maximum of 50 randomly selected SRAs for each of the 85 genera. These are listed in the supplementary file `sras.tab` The number of candidate reference genomes to be considered for each genus was capped at 200; the median number of genomes per genus was 9 (see Figure S3). The only genus with more than 200 reference genomes available was *Bordetella*. A maximum Mash distance was set to 0.1 (Ondov et al. 2016) between a preliminary assembly and a reference genome, as this was shown to be the maximum distance between the reference and sequenced isolate that riboSeed performs well with (see Waters et al. (2018) Figure 5). Run scripts are available in supplementary data; the reference genomes considered can be found in Supplementary file `reference_genomes.tab`.

### Benchmarking Re-assembled 16S rRNA against Complete Genome 16S rRNA sequences

sraFind was used to identify which SRA accessions corresponded to complete NCBI genomes for the genera considered in this study. These were matched with SILVA sequences sourced from complete genomes (see supplementary data “Provenance of strains”). Pairwise alignments were generated between the riboSeed 16S rRNA sequences and the SILVA sequences using the Biostrings package (Pagès et al. 2020) in “overlap” mode (a global alignment with free ends) with a simple scoring matrix (matches=1, mismatches=0); the highest-scoring alignment for each given reference 16S rRNA was used to identify misassemblies relative to the complete genome’s 16S rRNA sequence. Alignments shorter than 1400bp were rejected.

### Benchmarking Re-assembled 16S rRNA against Draft 16S rRNA sequences

Similar to the comparison to complete genomes above, we identified the SILVA sequences sourced from draft genome assemblies (see supplementary data “Provenance of strains”). Pairwise alignments were generated between the riboSeed 16S rRNA sequences and the SILVA sequences using the Biostrings package (Pagès et al. 2020) in “overlap” mode (a global alignment with free ends) with a simple scoring matrix (matches=1, mismatches=0); the highest-scoring alignment for each given reference 16S rRNA was used to identify missassemblies relative to the complete genome’s 16S rRNA sequence. Alignments shorter than 1400bp were rejected.

### Assessing Taxonomic Assignment

The DADA2 pipeline was used to process each of the four datasets in Table 1. The resulting sequence tables were combined, and we assigned taxonomy with the naive Bayes classifier implemented in DADA2. This classified sequences at the genus level, and DADA2’s “assignSpecies” command was used to assign species-level taxonomy; we enabled the “allowMultiple” parameter to view ambiguities in the assignment. This analysis was used to compare assignment with SILVA 132 alone and assignment with SILVA 132 augmented with sequences generated by Focus16. All scripts can be found in supplementary materials.

## Results

### Benchmarking Re-assembly Accuracy

riboSeed has been shown to generate high-quality reconstructions of each rDNA region when benchmarked against hybrid assemblies (Waters et al. 2018). Using sraFind, we identified which sequences in SILVA originated from closed, complete genomes; those genome with short-read SRAs were used to benchmark the accuracy of the 16S rRNA sequences re-assembled with the Focus16 pipeline (as described in Methods section) against the sequence in SILVA. In our dataset, 61 of these SRA/complete genome pairs were present.

Comparing the re-assembled 16S rRNA sequences to the 16S rRNA sequences from complete genomes shows a low error rate, with 53% of sequences being perfect reconstructions and 95% of sequences having fewer than 5 errors per Kbp (Figure 3). This confirms that Focus16’s best-case accuracy yields perfect reconstructions of the rDNA region; those cases for which reconstruction was imperfect rarely have more than 10 errors (an error rate rarely exceeding 0.7%), and 99% of sequences had fewer than 10 errors per Kbp. This suggests that sequences could be used to augment existing databases; the benefits and consequences of this are presented in the discussion. The error rates for amplicon data in SILVA are difficult to determine; under optimal conditions Sanger sequencing has very low error rates (Shendure and Ji 2008); however when multiple sequences are inadvertently sequenced at the same time (i.e. multiple copies from a single organism), the trace will reflect the differences as short, imperfect, or overlapping peaks. As the trace/quality data for amplicon sequences are not typically available, it is impossible to determine the accuracy of such sequences.

**Figure 3:**
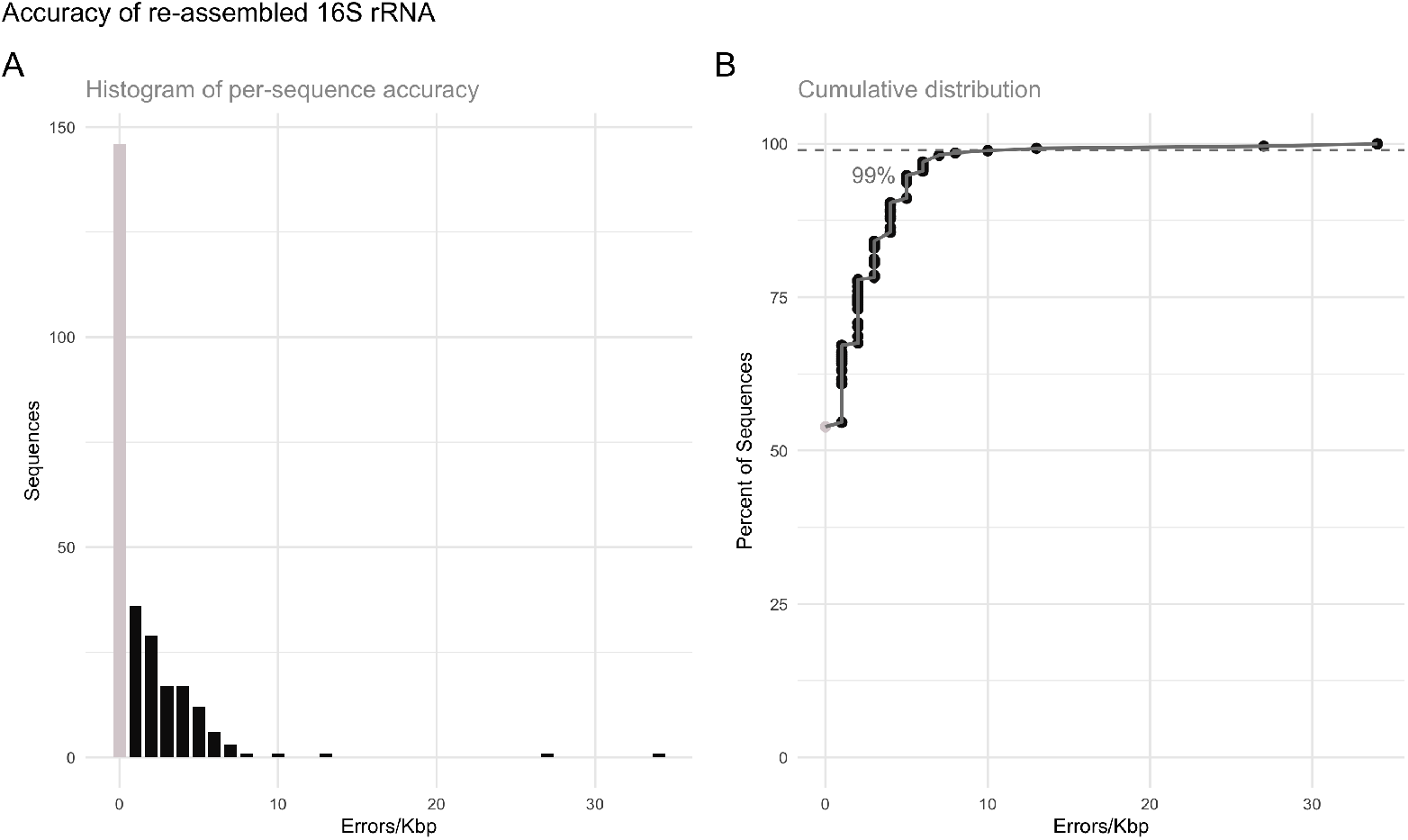
Assembly errors per kilobase calculated between each Focus16 sequence and the reference genome as counts (A) and cumulatively (B). In the genera considered, 285 16S rRNA sequences from 61 complete genomes were present in SILVA; riboSeed recovered 271 of these. 146 of these 16S rRNA alleles were identical between riboSeed and complete genome (grey bars).

### Comparing re-assembled 16S rRNA to draft 16S rRNA

As repeated rDNA operons are difficult to resolve with short read sequences, draft genome assemblies can (and often do) contain a single assembled rDNA region with elevated read coverage. This can be problematic for genus or 16S rRNA classification as the 16S rRNA recovered may not just correspond to one of several 16S rRNA copies, but it can be a consensus “summary”/“collapsed” 16S rRNA resulting from imperfect assembly of the repeated region. We provide a few examples of such alignments in Figure 4 (see all alignments in supplementary repository folder `figures/draft_alignments/`).

**Figure 4:**
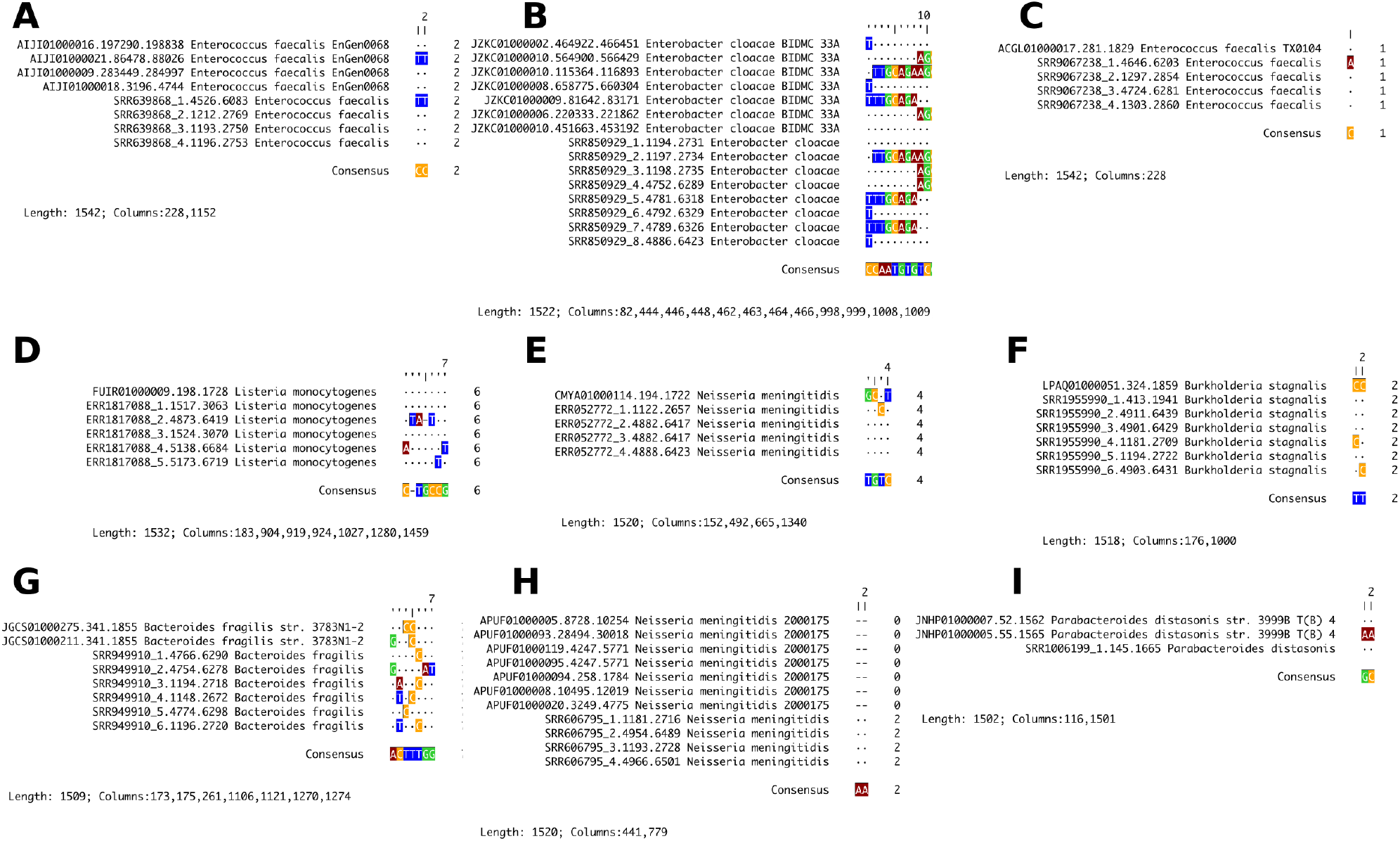
Representative SNP alignments comparing 16S rRNA sequences from original draft assemblies to the re-assembled sequences. Alignments were generated with DECIPHER and all columns matching the consensus were removed; original alignment length and column numbers for each SNP are shown under the sequence names. Names starting with an SRA accession such as ERR and SRR are the re-assembled sequences. The following types of relationships occur: all alleles recovered in original and re-assembled (A,B), sole original sequence misses a single (C) or multiple SNPs (D), disagreement between sole original and re-assembled alleles (E), original sequences appears to be amalgamation of alleles (FG), a deletion is present in original allele (H), re-assembly fails to reconstruct an allele (I).

In such cases, without the capacity to verify the regions with Sanger or long-read sequencing, determining which sequences are the missassemblies and which should be regarded as true is an impossible task.

### Augmenting SILVA with results from Focus16

#### Recovering Sequences from Re-assembly

Focus16 was used to build an extended database for the three mock datasets described in the DADA2 paper and a real-world dataset from the Endobiota study. From the 85 genera considered, Focus16 processed 2387 SRAs, and recovered 16S rRNA sequences from 1392 SRAs. The average execution time for a given SRA was approximately 23 minutes. Several factors can contribute to failing to recover 16S rRNA sequences from a given SRA, and among these are a too-distant reference genome, low rDNA flanking diversity, low read length, or high read error rates. In total, we recovered 5854 16S rRNA sequences, of which 3008 were unique.

#### Recovery of unique sequences

Ideally, Focus16 would be applied to every eligible SRA currently available, and periodically rerun as more high-quality reference genomes are generated with long-read technologies; in this pilot study, we assessed the increase in unique sequences gained by augmenting SILVA with only the 85 genera found across these four datasets. For thoroughly-sequenced genera such as *Escherichia*, *Pseudomonas*, or *Bacillus*, the increases in unique sequences are small. However, other taxa showed marked increases in the genus-level 16S rRNA sequence diversity (Figure 5).

**Figure 5:**
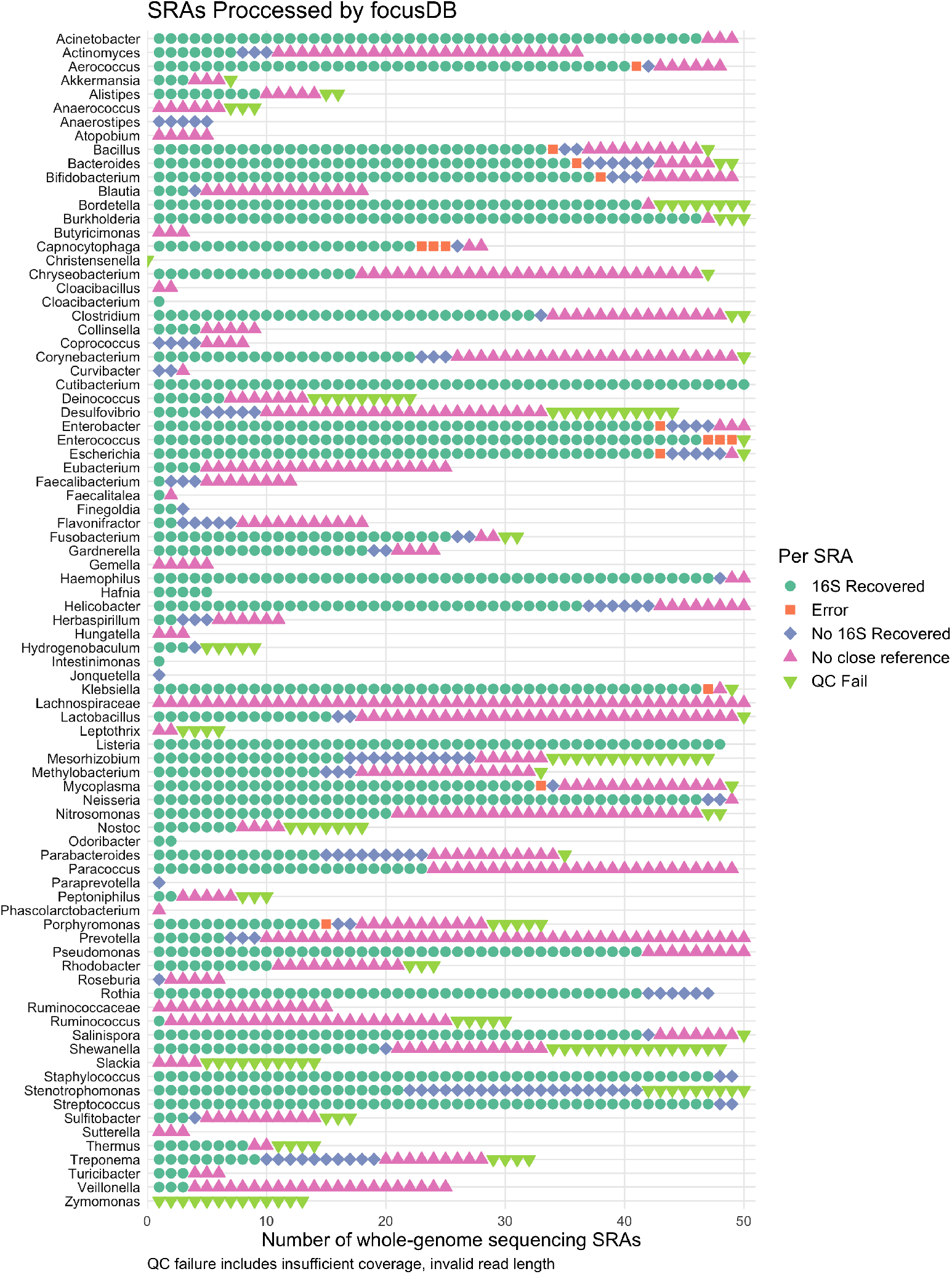
Summary of Focus16 outcomes for reassembling SRA datasets. Green circles indicate SRAs that yielded 16S rRNA sequences, while blue diamonds indicate SRAs failing to yield re-assembled 16S rRNA sequences. Pink triangles show SRAs that were rejected due to limitations in the diversity of available reference genomes, and inverted green triangles show SRAs rejected due to read length, insufficient coverage, poor read quality, etc. A few errors occurred, usually when the SRAs metadata conflicted with the associated sequencing data and caused download errors or errors from reads with incorrect pairing. In these cases, the datasets were discarded.

**Figure 6:**
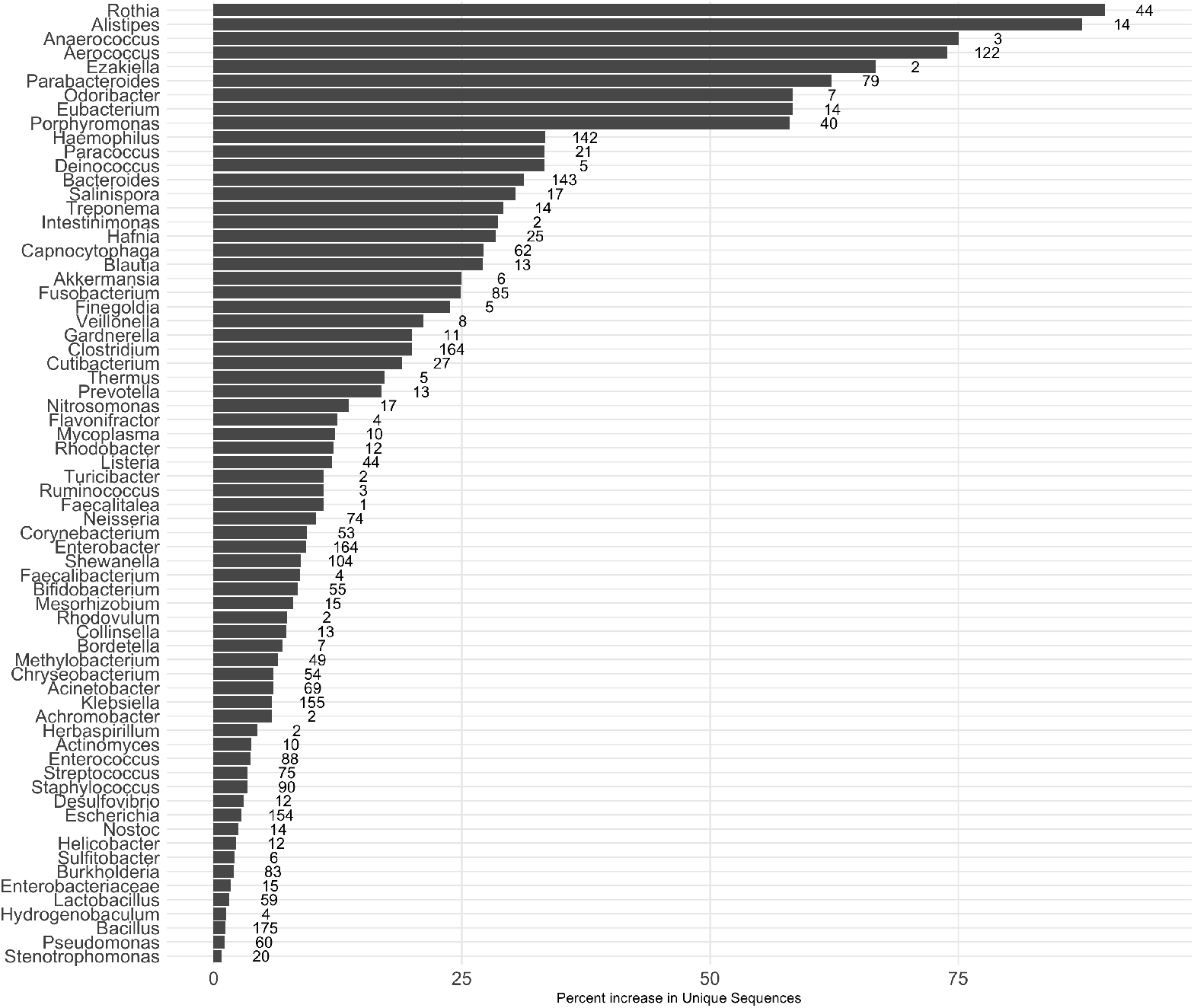
Percentage increase in unique sequences (and number of added unique sequences), by genus. The addition of the sequences recovered by Focus16 increases the number of unique sequences for the given taxa; the greatest increases are found in “under-sequenced” taxa and/or taxa with less well-conserved rRNA sequences.

### Assessing Taxonomic Assignment

DADA2 was used to identify ASVs from four datasets (Table 1), resulting in a total of 4098 ASVs (109 sequences from the “HMP” dataset, 26 from the “Extreme” dataset, 94 from the “balanced” dataset, and the rest from the EndoBiota study). We then compared the taxonomic results of classification using the SILVA 132 database either as-is, or augmented with novel 16S identified with Focus16.

Of the 4098 sequences, Focus16 changed the taxonomic assignment of 20 strains (see Table 2, or Supplementary file STABLE_different_assignment.tab for the actual sequences). Changes could happen in three ways: a previously unclassified ASV gained classification, a previously-assigned ASV gained more species- (or genus-) level details, or a previously-classified ASV was given a different classification. In our dataset, three unclassified strains gained annotations (Table 2 rows 5, 10, and 11). The remaining 17 had more detail added to the genus or species level; usually, this meant that with SILVA alone a single species classification was given, but with the augmented database, it was indicated that the ASV was ambiguous and could belong to more than one species (Table 2 rows 4,6,8,9,12,13,14) or genus (Table 2 rows 1,2,7, 15-20) level. No previously-assigned ASVs were assigned a completely different annotation.

**Table 2:** Of the 4098 ASVs aggregated across the four datasets considered, the augmented database modified taxonomic assignment of 20 sequences. Cells reading ‘NA NA’ indicate that no genus or species could be assigned. Cells reading NA speciesA/speciesB indicate that the lowest common ancestor for a sequence could not be determined at the genus level; in such cases DADA2 gives the designation NA rather than listing possible genera as is done at species level.

|  | SILVA | SILVA + Focus16 |
| --- | --- | --- |
| 1 | Peptoniphilus asaccharolyticus/ grossensis/ harei | NA asaccharolyticus/ grossensis/ harei/ massiliensis |
| 2 | Peptoniphilus harei | NA harei/ mediterraneensis |
| 3 | NA asburiae/ cancerogenus/ cloacae/ hormaechei/ ludwigii/ pneumoniae/ quasipneumoniae/ xiangfangensis | NA asburiae/ bugandensis/ cancerogenus/ cloacae/ hormaechei/ ludwigii/ pneumoniae/ quasipneumoniae/ xiangfangensis |
| 4 | Enterobacter asburiae/ cloacae/ hormaechei/ ludwigii/ mori/ soli/ tabaci/ xiangfangensis | Enterobacter asburiae/ bugandensis/ cloacae/ hormaechei/ ludwigii/ mori/ soli/ tabaci/ xiangfangensis |
| 5 | NA NA | Rothia mucilaginosa |
| 6 | Blautia massiliensis | Blautia hansenii/ massiliensis |
| 7 | Phascolarctobacterium faecium | NA faecium/ fermentans |
| 8 | Veillonella dispar | Veillonella dispar/ parvula |
| 9 | Gemella haemolysans/ sanguinis/ taiwanensis | Gemella haemolysans/ morbillorum/ sanguinis/ taiwanensis |
| 10 | NA NA | Haemophilus parainfluenzae |
| 11 | NA NA | Haemophilus parainfluenzae |
| 12 | Bacteroides uniformis | Bacteroides helcogenes/ uniformis |
| 13 | Alistipes massiliensis | Alistipes massiliensis/ shahii |
| 14 | Alistipes obesi | Alistipes finegoldii/ obesi/ shahii |
| 15 | Cloacibacterium normanense | NA normanense/ taklimakanense |
| 16 | Phascolarctobacterium faecium | NA faecium/ fermentans |
| 17 | Listeria innocua/ ivanovii/ marthii/ monocytogenes/ phage/ seeligeri/ welshimeri | NA epidermidis/ innocua/ ivanovii/ marthii/ monocytogenes/ phage/ seeligeri/ welshimeri |
| 18 | Listeria innocua/ ivanovii/ marthii/ monocytogenes/ phage/ seeligeri/ welshimeri | NA epidermidis/ innocua/ ivanovii/ marthii/ monocytogenes/ phage/ seeligeri/ welshimeri |
| 19 | Rhodobacter johrii/ megalophilus/ ovatus/ sphaeroides | NA johrii/ megalophilus/ ovatus/ sphaeroides/ sulfidophilum |
| 20 | Rhodobacter johrii/ megalophilus/ ovatus/ sphaeroides | NA johrii/ megalophilus/ ovatus/ sphaeroides/ sulfidophilum |

If this pilot study is perfectly representative, users could expect an improved taxonomic assignment for about 0.5% of ASVs; two factors must be considered before extrapolating that value beyond this study. First, we were limited in the number of SRAs per genus that could be processed (Supplementary Figure 1 shows the number of SRAs per genus). Second, and perhaps more importantly, the majority of genera in this study are associated with human microbiomes, an area which has already seen an extensive amount of focus in terms of amplicon sequencing, whole-genome sequencing, and genome completion efforts. Other environments have not had this benefit, and perhaps have more room for improvement.

## Discussion

Focus16 orchestrates the re-assembly of whole-genome sequencing datasets in SRA to recover 16S rRNA sequences that may be missing from the existing reference databases. Using riboSeed, Focus16 re-assembles draft genomes that currently contribute a single (often collapsed) 16S rRNA to resolve distinct instances of the 16S rRNA operon. We show that this increases the sequence diversity (number of unique sequences) of the 16S rRNA databases, and that the increased diversity results in measurable improvements to taxonomic assignment.

Focus16 improved fine-scale taxonomic assignment in two ways: by assigning previously unclassified sequences, and by revealing “overeager” species assignment when a 16S rDNA sequence could have come from two or more species. While at face value this appears to reduce the precision of taxonomic assignment, it reveals cases where species-level assignment was inappropriate. Based on the improved taxonomic assignment in this pilot study of 85 genera, we believe a wide-scale application of Focus16 could benefit the community.

A natural concern about the approach presented here is the danger of “poisoning” the database with sequences that may or may not be 100% accurate. This is valid concern, but one we believe to be outweighed by the potential of offsetting the known problems with existing 16S rRNA databases. Those 16S rRNA sequences in SILVA originating from amplicons lack to the taxonomic confidence that comes with whole-genome sequencing. Sequences from draft whole-genome assemblies have known issues with rDNA missassembly; in the best case, only one accurate 16S rRNA is represented; in the worst case, the one assembled 16S rRNA sequence may be an amalgamation of the different copies. Until long-read sequencing efforts sufficiently explore the same microbial genomic diversity currently covered by current 16S rRNA databases, these issues must be considered when attempting community analysis via 16S rRNA sequencing. A conservative approach to utilizing sequences recovered with the methodology presented here may be to incorporate a measure of taxonomic assignment confidence, where references sequences originating from long reads, amplicons, draft assemblies, and focus16 reassemblies could be weighted appropriately.

However, there are three main limitations facing the large-scale application of Focus16: the first is the bandwidth, computational power, memory, and storage required to re-assemble the 98,329 SRAs (as of October 2019) that were used to generate draft genomes. Given sufficient storage and unfettered access to a medium/large computing cluster such as one supporting a university or research institute (say 150 compute nodes), the task could be completed within two weeks^1^, and the task could be accomplished in even less time with a sufficiently large cloud computing budget; however with modest hardware (8 cores, 20GB RAM), this would take about 4 years in “–fast” mode. These estimates are ignoring the 112,695 draft genomes for which no reads were ever released, which leads to the second limitation: data availability. Releasing draft genomes without the reads used to generate them hampers efforts such as this one to expand beyond the purpose of the original study.

The third limitation of this approach is the availability of high-quality closed genomes to use as references. With the increased adoption of long read technologies, we envisage that this limitation will decrease with time; re-running the pipeline as new, complete reference genomes are generated will allow for ongoing improvements to the databases. Eventually, a point will come when Focus16 will no longer be needed as all candidate SRAs have been sufficiently utilized.

Further limitations exist within Focus16 and within riboSeed. The success of riboSeed’s *de fere novo* assembly hinges on the similarity of the reference to the sequenced isolate, the differentiating power of the rDNA flanking regions, read length, and other factors. This is one reason why not all SRAs yielded perfect rDNAs. Additionally, riboSeed does not currently support mate-paired libraries; these are much less widely used than the typical single-end or paired-end libraries used in short-read sequencing.

Despite these limitations, we have shown that Focus16 can contribute towards better molecular ecology analysis; augmenting SILVA with the sequences re-assembled from the 85 genera considered here led to a small increase in the number of unique sequences in the database. Using the augmented database for taxonomic assignment revealed some limitations of low-level taxonomic assignment, and led to the classification of additional ASVs. We invite the community to consider augmenting existing databases (such as NCBI’s 16S RefSeq Microbial database, SILVA, RDP, and GreenGenes) with the approach outlined here.

## Acknowledgements

Many thanks to Christopher Quince for the helpful conversations on the topic.

## Data Accessibility

The code for Focus16 can be found at https://github.com/FEMLab/focus16; the code for all the analyses presented in this work can be found at https://github.com/FEMLab/focus16_manuscript. All data used is archived at Zenodo accession 10.5281/zenodo.3956433.

## Author Contributions

Author contributions according to the CRediT taxonomy (Allen, O’Connell, and Kiermer 2019) are listed alphabetically as follows: Conceptualization: LP, NW ; Methodology: FA, BN, LP, NW; Software and Data Curation: BN, NW; Validation: BN, LP, NW; Formal analysis: BN, LP, NW; Investigation: BN, NW; Resources; FA, VOF, LP; Writing - Original Draft: BN, NW; Writing - Review & Editing: FA, FB, VOF, AH, BN, LP, NW; Visualization: BN, LP, NW; Supervision: FA, FB, VOF, AH, LP, NW; Project administration FA, VOF, NW; Funding acquisition: FA, FB, VOF, LP, NW.

## Footnotes

1 This estimation is based on an rough average processing time of 23 minutes per run, but this is highly dependent on download speed, read/write speed, genome size, and sequencing depth.

