## Supplementary Methods 1 for "Mining underutilized whole-genome sequencing projects to improve 16S rRNA databases"

### 1 Introduction

This document contains all the of steps needed to reproduce the findings presented in this work, with the exception of the community analyses described in `DADA2_analyses.Rmd`; it was written with literate programming, so all code can be viewed in the source document at https://github.com/femlab/focus16_manuscript/blob/master/docs/.

#### 1.1 Third-party software versions

The conda environment used for the analyses is defined in https://github.com/femlab/focus16_manuscript/blob/master/focus16_env.txt.

### 2 Data Availability

#### 2.1 Number of SRAs available for each species

A python script `sralist.py` was used to generate a list of the number of SRAs for a given species, based on parsing the results from sraFind from 2019-10-24. The input files for the “Balanced,” “HMP,” and “Extremes” were created from the descriptions of the mock communities found in the their respective publications(Schirmer et al. 2015; Kozich et al. 2013; Callahan et al. 2016; Ata et al. 2019). Generating the Endobiota species file is described in `DADA2_analyses.Rmd` section “Preliminary taxa assignment for EndoBiota study.”

The results per dataset are shown in Figure S1. Because some genera have thousands of SRAs, we limited subsequent analyses to 50 SRAs per genus.

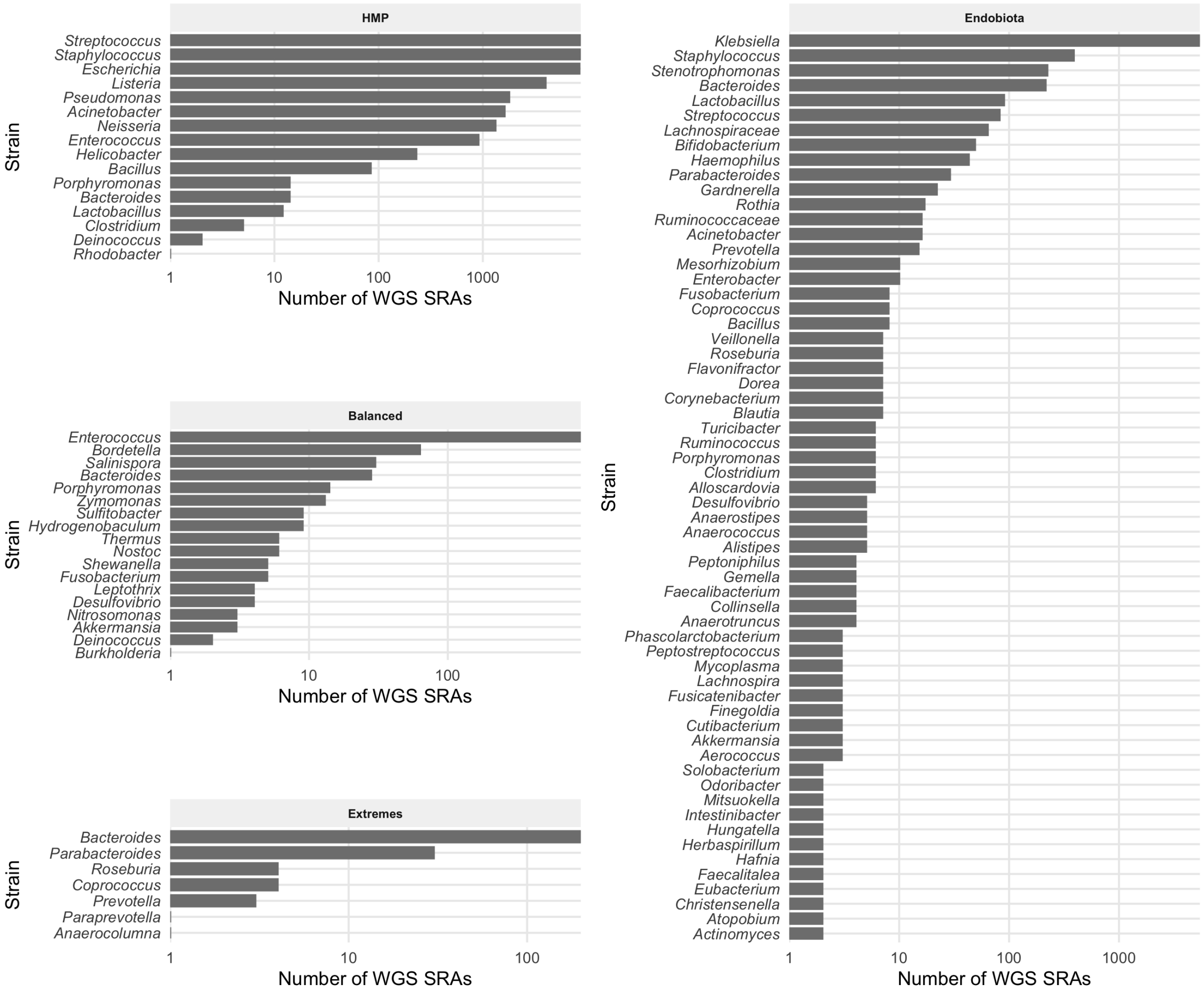

Figure S 1: Whole-genome sequencing SRA data availability for strains in the four datasets. Bars show the number of SRAs per genus as of 2019-10-24. Genera with no publicly available SRAs are removed; results are displayed on a log scale. 109 of 193 genera have available SRA data. Clinically-relevant genera tend to have many SRAs.

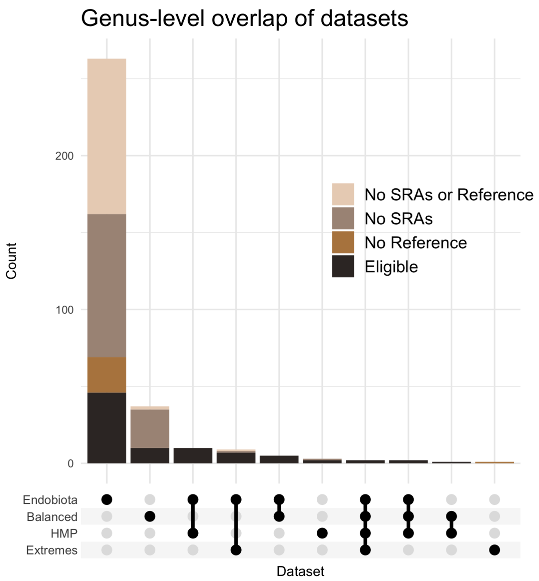

| | Eligible | No SRAs | No SRAs or Reference | No Reference |
|---|---|---|---|---|
| *Balanced* | 20 | 25 | 2 | 0 |
| *Endobiota* | 72 | 94 | 102 | 23 |
| *Extremes* | 9 | 1 | 1 | 1 |
| *HMP* | 17 | 1 | 0 | 0 |

Figure S 2: 368 genera across the four datasets were considered. Of the 333 unique genera, 85 met our criteria of having public short-read data in the SRA and at least one RefSeq complete genome available. The x axis shows the overlapping sets of genera among the four datasets; the color shows how many of the genera met the inclusion criteria. While only a fraction of the SRAs meet the requirements for reassembly, future data availability will allow us to revisit previously-excluded SRAs.

### 3 Running Focus16 on candidate taxa

#### 3.1 Parameter choice

Below is a copy of the script used to run the pipeline for the list of genera determined above. The following parameters were used:

- `--n_references 200`: the maximum number of RefSeq genomes to consider for use as a reference
- `--n_SRAs 50`: limits reassembly to 50 random SRA read sets per genus
- `--run_de_novo_control`: ensures riboSeed run a control *de novo* assembly with SPAdes. This is used for comparisons of the *de novo* assembly to the *de fere novo* assemblies.
- `--maxdist .1`: set the maximum Mash distance between a reference genome and a preliminary assembly
- `-v 1`: sets verbosity
- `--fastqtool fastq-dump`: prefer fastq-dump to fast(er)q-dump
- `--timeout 1500`: sets download limit for 25 minutes; we found downloads lasting long than that usually were hanging
- `--process_partial`: if a download times out, proceed with the reads downloaded
- `--use_available`: use any reads downloaded while previously processing an SRA
- `--sge`: distribute assembly tasks as an SGE array job on the HPC
- `--sge_env 16db`: name of the conda environment that the distributed jobs to be executed with
- `--njobs 6` : number of tasks to be run concurrently for each SGE array
- `--threads 2`: resource allocation for main job
- `--cores 6`: resource allocation for main job
- `--memory 20`: resource allocation for main job

```bash
#!/bin/bash
#$ -t 1-85
#$ -tc 15
#$ -cwd
#$ -j yes
#$ -N comb_genus
#$ -pe mpi 4
#$ -l h_vmem=20G
set -e
counter=1

cd /mnt/shared/scratch/synology/nw42839/2019-11-04-focusdb/
while read genus species
do
    if [ "$counter" -eq "$SGE_TASK_ID" ]
    then
        conda activate 16db
        focusDB  -o ${genus}_genus --organism_name "${genus}" --njobs 6 --threads 2 \
        --cores 6 --memory 20 --n_references 200 --n_SRAs 50 --maxdist .1 \
        --focusDB_data /mnt/shared/scratch/nw42839/.focusDB/ -v 1 --timeout 1500 \
        --fastqtool fastq-dump --process_partial --use_available --sge --sge_env 16db
    fi
    counter=$((counter + 1))
done < docs/datasets/combined_genuses.txt
```

Afterwards, the results were aggregated, and then were moved to this repo under the `results` dir.

```
mkdir 2019-12-19-results
while read genus; do echo $genus; cp ./${genus}_genus/${genus}_fast_ribo16s.fasta ./2019-12-19-results/
cp --parents ./*_genus/*/results/riboSeed/seed/final_de_novo_assembly/contigs.fasta ./2019-12-19-result
cat ./*_genus/SUMMARY > 2019-12-19-results/summary_all
cat ./2019-12-19-results/*seq_summary.tab > 2019-12-19-results/sequence_summary_all.tab
tar czf 2019-12-19-results.tar.gz 2019-12-19-results/
```

##### 3.2 List of References Considered

After execution of the pipeline, all of the references considered were recorded as follows:

```
while read genus ; do \
for j in  .focusDB/references/$genus/*.fna; do \
echo -e "${genus}\t${j}" >> docs/reference_genomes.tab ; \
done ;done < docs/datasets/combined_genuses.txt
```

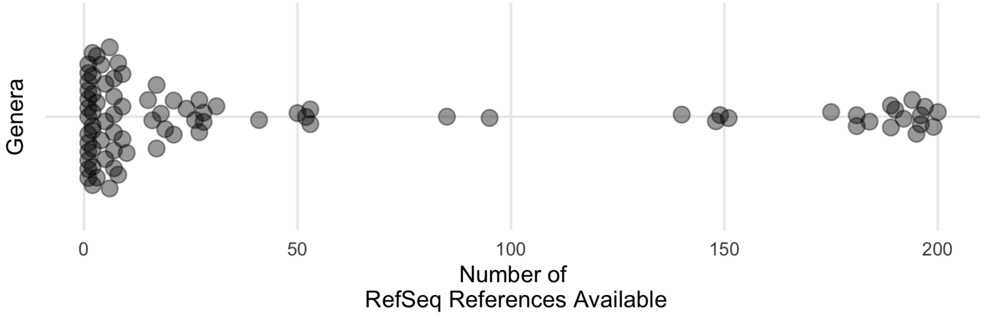

Figure S 3: A median of 9 references were available for each of the genera considered. The number of references per genera was capped at 200. In practice, this only effected the Bordetella genus. Data were plotted with ggbeeswarm using the quasirandom layout to show the distribution.

##### 3.3 List of SRAs

After the pipeline was run, the SRAs processed for each genus was recorded.

```
ls -d results/2019-12-19-results/*_genus/* > docs/sras.tab
```

#### 4 Quality Control

##### 4.1 Diagnosing poor assemblies

In the course of comparing the reassembled sequences to those for which reference genomes were available, we determined that the pipeline struggled in two cases: the *Treponema* genus, and one *Bacillus anthracis* SRA: SRR2155541. We do not currently know the issue with the *Treponema* genus, but using blobtools we determined that SRR2155541 is likely contaminated with *Micrococcus luteus* (Figure 4).

```
# we downloaded GCF_000008165.1_ASM816v1_genomic.fna as a reference
docker run --memory 16G --rm  -v $PWD:/home/ nickp60/ezblobtools \
  -r /home/results/2019-12-19-results/Bacillus_genus/SRR2155541/results/riboSeed/seed/final_de_novo_ass
  -d ref_prok_rep_genomes -o /home/GitHub/focusdb/docs/blob_SRR2155541/ -t 2 -m 16
```

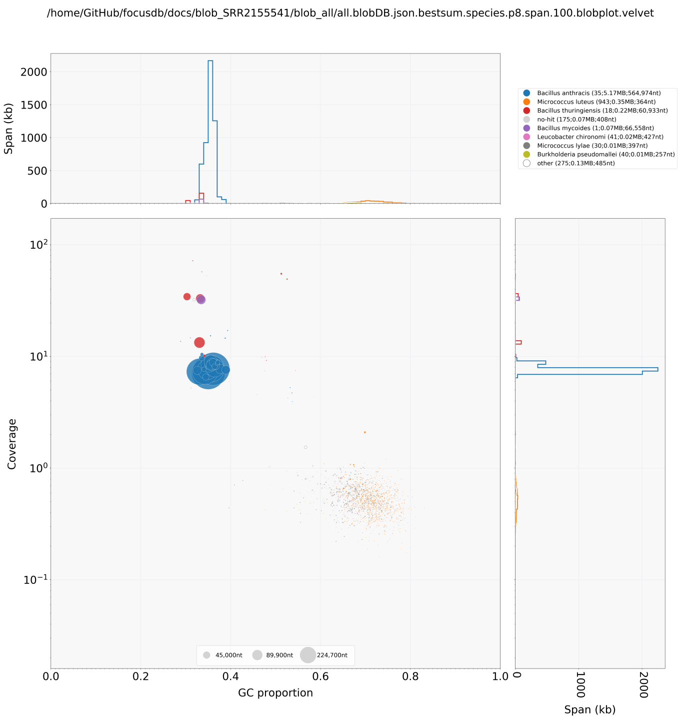

Figure S 4: Assessing the contents of *Bacillus anthracis* SRA SRR2155541 showed possible contamination with *Micrococcus luteus*, as can be seen from the coverage and GC content differences.

##### 4.2 Identifying poor taxonomic assignments

In addition to those mentioned above with poor assemblies, we filtered out any SRAs with inconclusive domain-level agreement, where Kraken reports the top level Domain taxonomy with less than 70% agreement . This primarily intended to remove datasets with poor host-DNA depletion.

Figure 5 shows a histogram of the domain-level agreement. We parse the outputs from Kraken2's taxonomic assignment, and flag any with a low (<70) percent of domain-assigned sequences, as this could indicate that an SRA is contaminated. Some (such as Treponema SRAs SRR3571775 and SRR3584844) have $\approx$ 60% human DNA, indicating incomplete removal of host DNA, and as such we drop them, losing 9 SRAs. However, it should be noted that BLASTing contigs from these assemblies appear to show bacterial DNA – the Kraken2 database itself may have contamination where Treponema kmers are labelled as human. Either way, these (and any other under the 70% domain-level assignment threshold) are dropped.

#### 5 Results

##### 5.1 Assessing per-SRA outcomes of the pipeline

The outcome of each SRA was determined after running the pipeline. For the purpose of this figure, the following messages were combined:

- `Error`:
  - Any errors involving riboSeed
  - Error downloading datasets or references
  - Unknown errors
- `QC Fail`:
  - Invalid library type (eg mate-paired reads)

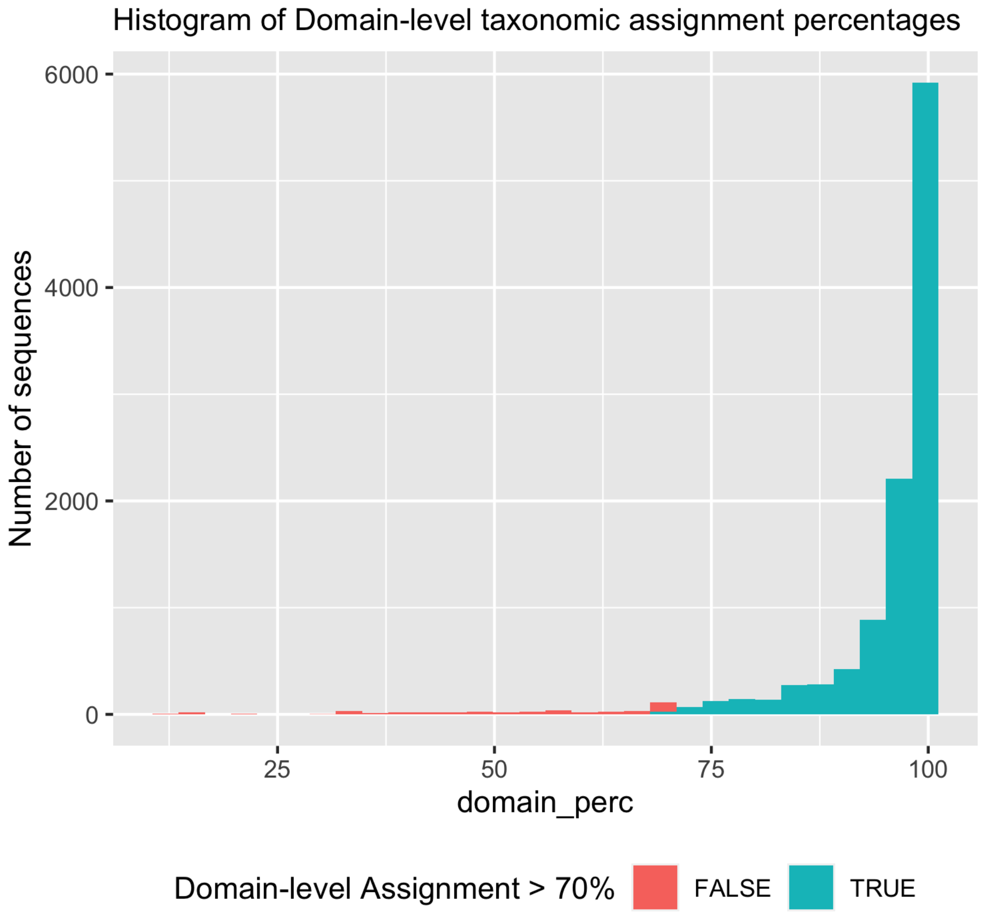

Figure S 5: Datasets where Kraken's taxonomic assignment of preliminary assembly contigs showed less less than 70 percent agreement were removed, as these are likely contaminated with host DNA

- Reads < 65bp (eg early Genome Analyzer reads)
- Reads > 300bp (eg PacBio or Nanopore reads)
- Insufficient coverage < 15x

#### 5.2 Extracting 16S rRNA from de novo assemblies

For the comparison of error rates between the *de fere novo* assembly and *de novo* assembly, `make_silva_style_db_from_contigs.py` was used to identify 16S rRNA regions using Barrnap and extract those regions from the *de novo* contigs .

#### 5.3 Assessing uniqueness

First, for simplicity, we combine all the sequences for all the runs. We have the logs to relate the sequences to why they were selected, but as we are talking about improving the database as a whole, combining them all simplifies aspects of interpretation.

We removed any from the list as identified above in section "Identifying poor taxonomic assignments."

#### 5.4 Assessing SILVA's composition

We read in the SILVA database, and parsed the headers using the prefix mapping found at https://www.ncbi.nlm.nih.gov/Sequin/acc.html. Each header for the SILVA database consists of `>ACCESSION.start.end name`. This revealed the origin of the individual sequences in the database.

#### 5.5 Assessing the provenance of the SILVA strains

16S sequences in databases such a SILVA, GreenGenes,or RDP can come from several sources, namely: - complete genomes (in RefSeq, Genbank, etc) - draft genomes (in the NCBI's Assembly database) - 16s amplicons from Sanger sequencing - 16s amplicons from high-throughput amplicon sequencing.

As repeated rDNA operons are difficult to assembly, draft genomes can (and often do) contain a single rDNA. This is problematic for species ideentification – the rDNA recovered is not just one of $n$ rDNAs, but it can be a consensus "summary" rDNA resulting from trying to assemble the repeated region.

riboSeed has been show to generate high-quality reconstructions of each rDNA when benchmarked against hybrid assemblies. Here, we compare the 16s sequences from riboSeed reassembly of draft genomes to the initial (potentially collapsed) 16s sequences.

As of release 0.8, the WGS master record is a column present as "WGS" in sraFind.tab. This allows us to relate the sequences in the SILVA headers to their original SRAs, so we can compare riboSeed's assemblies to those for which a complete genome is available, and assess the types of errors/collapses we see where only draft genomes are available.

Next we subsetted the dataframe to include only those that had sras from this study,(and by proxy, those that were sequenced with illumina)

Now we have identified which SRAs relate to which assemblies, and which we have analyzed. The first order of business is to compare the accuracy of fast mode (using riboSeed's `--just_seed` parameter) to 16S's from full assemblies. However, for a full analysis, we also need to compare to the *de novo* assembly (which we enabled using the `--run_de_novo_control` flag). Interactive outputs are created as well.

phantomjs was used to convert html alignments into images. These were selected to show the types of arrangements that occur comparing draft to the reassembled genomes. Inkscape was used to combine images.

#### 5.6 Calculating total SRAs needing re-assembly

Over the course of the development of Focus16 we estimated the run-time for a subset of SRAs; runtimes for the whole dataset were difficult to determine due to the time stamps not reflecting I/O or network limitations.

This analysis calculates run time estimates based on an early run of the genera in the HMP dataset.

Extrapolating time of execution for the entire SRA was based off the 98329 SRAs available as of October 2019, the last update of sraFind.

Ata, Baris, Sule Yildiz, Engin Turkgeldi, Vicente Pérez Brocal, Ener Cagri Dinleyici, Andrés Moya, and Bulent Urman. 2019. "The Endobiota Study: Comparison of Vaginal, Cervical and Gut Microbiota Between Women with Stage 3/4 Endometriosis and Healthy Controls." *Scientific Reports* 9 (1): 1–9. https://doi.org/10.1038/s41598-019-39700-6.

Callahan, Benjamin J., Paul J. McMurdie, Michael J. Rosen, Andrew W. Han, Amy Jo A. Johnson, and Susan P. Holmes. 2016. "Dada2: High-Resolution Sample Inference from Illumina Amplicon Data." *Nature Methods* 13 (7): 581–83. https://doi.org/10.1038/nmeth.3869.

Kozich, James J., Sarah L. Westcott, Nielson T. Baxter, Sarah K. Highlander, and Patrick D. Schloss. 2013. "Development of a Dual-Index Sequencing Strategy and Curation Pipeline for Analyzing Amplicon Sequence Data on the MiSeq Illumina Sequencing Platform." *Applied and Environmental Microbiology* 79 (17): 5112–20. https://doi.org/10.1128/AEM.01043-13.

Schirmer, Melanie, Umer Z. Ijaz, Rosalinda D'Amore, Neil Hall, William T. Sloan, and Christopher Quince. 2015. "Insight into Biases and Sequencing Errors for Amplicon Sequencing with the Illumina MiSeq Platform." *Nucleic Acids Research* 43 (6): e37–37. https://doi.org/10.1093/nar/gku1341.
