## Supplementary Methods 2 for "Mining underutilized whole-genome sequencing projects to improve 16S rRNA databases"

*last update: 01 April, 2020*

#### Contents

**1 Overview** **1**

**2 Endobiota study** **1**
2.0.1 Preliminary taxa assignment for Endobiota study . . . . . . . . . . . . . . . . . . . . . . 3

**3 Extremes dataset** **4**

**4 Balanced Dataset** **5**

**5 HMP** **5**

**6 Combining sequence tables** **6**

**7 Recreating the DADA2-formatted SILVA and augmented databases** **7**

**8 Assigning taxonomy to the combined sequence table** **7**

### 1 Overview

Here, we assess the effect of using the augmented dataset, we show the taxonomic assignment for 4 dataset: the "Balanced", "HMP", and "Extreme" dataset assessed with DADA2, and data from the recent "Endobiota" study.

The preprocessing of all the data is preformed in the next four sections, and then the final section describes combining all the sequences into one sequence table matrix for simplicity of manipulation of the results, while keeping track of which sequence comes from where.

DADA2 is actively being improved; because of this, we used the guidelines outlined in the tutorial rather than those originally described for these analysis in the DADA2 supplementary material.

### 2 Endobiota study

We follow a generic DADA2 workflow for the Endobiota BioProject PRJEB26800. Plots describing the quality of the read sets can be found in "end_R_q.png" and "end_F_q.png". Sample ERR2586009 was removed due to low sequence depth and poor quality.

```
# go to https://www.ncbi.nlm.nih.gov/sra?linkname=bioproject_sra_all&from_uid=572651
# hit send to run selector
#  Click to download "Run Accessions"
while read x; do fasterq-dump --split-files $x ; done < ~/Downloads/SRR_Acc_List-1.txt
```

```
filtpath <- "./docs/microbiome_data/clean/"
mb_meta <- read.csv2("./docs/microbiome_data/metadata.tsv", stringsAsFactors = F,sep="\t",comment.char = '#')
sra_meta <- read.csv("./docs/microbiome_data/SraRunTable-PRJEB26800.txt", stringsAsFactors = F,sep=",")
meta <- full_join(
  mb_meta %>%select(sample_alias, sample_description),
  sra_meta %>% select(Run, Sample.Name), by=c("sample_alias"="Sample.Name")
```

```r
)
meta$site <- gsub("(.*?) .*", "\\1", meta$sample_description)
meta$status <- gsub(".*\\((.*?)\\)$", "\\1", meta$sample_description)

# Forward and reverse fastq filenames have format: SAMPLENAME_R1_001.fastq and SAMPLENAME_R2_001.fastq
endo_fnFs <- sort(list.files(dirpath, pattern="_1.fastq.gz", full.names = TRUE))
endo_fnRs <- sort(list.files(dirpath, pattern="_2.fastq.gz", full.names = TRUE))
# Extract sample names, assuming filenames have format: SAMPLENAME_XXX.fastq
sample.names.endo <- sapply(strsplit(basename(endo_fnRs), "_"), `[`, 1)
```

```r
ggsave(
  plotQualityProfile(endo_fnFs),
  filename = file.path(dirpath, "end_F_q.png")    ,
  dpi = 300, width = 16, height = 10, units = "in")
ggsave(
  plotQualityProfile(endo_fnRs),
  filename = file.path(dirpath, "end_R_q.png")    ,
  dpi = 300, width = 16, height = 10, units = "in")

# We noticed that one of the datasets should be removed,
# as the sequencing quality appears severly impaired
baddiesF <-c(sample.names.endo[grep("009",sample.names.endo)])
baddiesR <-c(sample.names.endo[grep("009",sample.names.endo)])

sample.names.endo <- sample.names.endo[!sample.names.endo %in% unique(c(baddiesF, baddiesR))]
for (baddie in c(baddiesF, baddiesR)){
  endo_fnFs <- endo_fnFs[!grepl(baddie, endo_fnFs)]
  endo_fnRs <- endo_fnRs[!grepl(baddie, endo_fnRs)]
}
# Place filtered files in filtered/ subdirectory
endo_filtFs <- file.path(dirpath, "filtered", paste0(sample.names.endo, "_F_filt.fastq.gz"))
endo_filtRs <- file.path(dirpath, "filtered", paste0(sample.names.endo, "_R_filt.fastq.gz"))
names(endo_filtFs) <- sample.names.endo
names(endo_filtRs) <- sample.names.endo


endo_out <- filterAndTrim(
  endo_fnFs, endo_filtFs,
  endo_fnRs, endo_filtRs,
  trimLeft=30, trimRight=40,
  maxN=0, maxEE=c(2,2), truncQ=2, rm.phix=TRUE,
  compress=TRUE, multithread=TRUE)


endo_errF <- learnErrors(endo_filtFs, multithread=TRUE)
endo_errR <- learnErrors(endo_filtRs, multithread=TRUE)
endo_derepF <- derepFastq(endo_filtFs, verbose=TRUE)
endo_derepR <- derepFastq(endo_filtRs, verbose=TRUE)

endo_dadaFs <- dada(endo_derepF, err=endo_errF, multithread=TRUE)
endo_dadaRs <- dada(endo_derepR, err=endo_errR, multithread=TRUE)

endo_merger <- mergePairs(endo_dadaFs, endo_derepF, endo_dadaRs, endo_derepR, verbose=TRUE)

endo_seqtab <- makeSequenceTable(endo_merger)
summary((nchar(getSequences(endo_seqtab))))
hist(nchar(colnames(endo_seqtab)))
```

```r
# here we trim some of those shorter sequences
endo_seqtab2 <- endo_seqtab[,nchar(colnames(endo_seqtab)) %in% 360:450]
hist(nchar(colnames(endo_seqtab2)))
dim(endo_seqtab2)
# [1]    83 23152
endo_seqtab.nochim <- removeBimeraDenovo(
  endo_seqtab2, method="consensus",
  multithread=TRUE, verbose=TRUE)
sum(endo_seqtab.nochim)/sum(endo_seqtab2)
# [1] 0.9577382
dim(endo_seqtab.nochim)
#  [1]   83 3876

getN <- function(x) sum(getUniques(x))
endo_track <- cbind(
  endo_out,
  sapply(endo_dadaFs, getN),
  sapply(endo_dadaRs, getN),
  sapply(endo_merger, getN),
  rowSums(endo_seqtab.nochim))
# If processing a single sample, remove the sapply calls: e.g. replace sapply(dadaFs, getN) with getN(dadaFs)
colnames(endo_track) <- c("input", "filtered", "denoisedF", "denoisedR", "merged", "nonchim")
rownames(endo_track) <- sample.names.endo
head(endo_track)

#save.image("./endo.RData")
save(endo_seqtab.nochim, file = "./endo_seqtab.RData")
```

##### 2.0.1 Preliminary taxa assignment for Endobiota study

To do the initial analysis for identifying the genera present in the Endobiota samples, the following commands were used to assign taxonomy:

```r
endo_taxa <- assignTaxonomy(
  endo_seqtab.nochim,
  "~/Downloads/silva_nr_v132_train_set.fa.gz", multithread=TRUE)
endo_taxa_species <- addSpecies(endo_taxa, "~/Downloads/silva_species_assignment_v132.fa.gz")
taxa.print <- endo_taxa_species # Removing sequence rownames for display only
taxa.print[is.na(taxa.print[,6])] <- ""
taxa.print[is.na(taxa.print[,7])] <- ""
thesenames <- gsub("^ $", "", paste(taxa.print[,6], taxa.print[,7] ))
table(thesenames)

write.table(
  sort(unique(thesenames)),
  file = "./docs/microbiome_data/endo_species.txt",
  row.names = F,  col.names = F, quote=F)
# here, if the genus level annotation is something like "group" or "clade" or
# #otherwise unhelpful, we tidy it up, or use the "family" level annotation
# if needed.  These get fixed later when making the combined list for the actual run.
fg <-  data.frame(endo_taxa_species[,c(5,6)], stringsAsFactors = F)
rownames(fg) <- NULL
fg$cleangenus <- gsub("[-_](.*)", "", fg$Genus)
fg$cleanfamily <- gsub("[-_](.*)", "", fg$Family)
fg$cleangenus <- ifelse(grepl("\\d", fg$cleangenus),fg$cleanfamily, fg$cleangenus)
fg$cleangenus <- ifelse(is.na(fg$cleangenus),fg$cleanfamily, fg$cleangenus)
# fix escherichia/shigella
# fg$cleangenus[gsub("(.*)\\/(.*)", "\\1", fg$cleangenus) !=  fg$cleangenus]
```

```
fg$cleangenus <-  gsub("(.*)\\/(.*)", "\\1", fg$cleangenus)
write.table(
  sort(unique(fg$cleangenus)),
  file = "./docs/microbiome_data/endo_genus.txt",
  row.names = F,  col.names = F, quote=F)
```

### 3 Extremes dataset

Here, we largely use the parameters defined in the original paper, but we updated to use the commands recommended in the tutorial for version 1.12.

```
extremes_path <-  "."
ex_fnF <- file.path(extremes_path, "SRR2990088_1.fastq")
ex_fnR <- file.path(extremes_path, "SRR2990088_2.fastq")

ggsave(
  plotQualityProfile(ex_fnF),
  filename = file.path(dirpath, "ex_F_q.png"),
  dpi = 300, width = 16, height = 10, units = "in")
ggsave(
  plotQualityProfile(ex_fnR),
  filename = file.path(dirpath, "ex_R_q.png"),
  dpi = 300, width = 16, height = 10, units = "in")
#  Forward reads are reasonably high quality.
#  Trimming the first 20 nts, and last 10 (truncate at 240).
# Reverse read quality drops off substantially.
# Trimming the first 20 nts, and last 50 (truncate at 200).

ex_filtF <- "ExtremeF_EE2.fastq.gz"
#ex_filtFO <- "ExtremeFO_EE2.fastq.gz"
ex_filtR <- "ExtremeR_EE2.fastq.gz"
ex_out <- filterAndTrim(
  fwd = ex_fnF, filt = ex_filtF,
  rev = ex_fnR, filt.rev = ex_filtR,
  maxN=0, maxEE=2, truncQ=2,
  truncLen=c(240,200), trimLeft=c(20,20), compress=TRUE, verbose=TRUE)
# Kept about 60 percent of the paired reads and
# 70 percent of the forward-only reads.

## Run DADA2 Pipeline
ex_errF <- learnErrors(ex_filtF, multithread=TRUE)
ex_errR <- learnErrors(ex_filtR, multithread=TRUE)
ex_dadaF <- dada(ex_filtF, err=ex_errF, multithread=TRUE)
ex_dadaR <- dada(ex_filtR, err=ex_errR, multithread=TRUE)
ex_mergers <- mergePairs(ex_dadaF, ex_filtF, ex_dadaR, ex_filtR, verbose=TRUE)

ex_seqtab <- makeSequenceTable(ex_mergers)

hist(nchar(colnames(ex_seqtab)))
ex_seqtab.nochim <- removeBimeraDenovo(
  ex_seqtab, method="consensus",
  multithread=TRUE, verbose=TRUE)
sum(ex_seqtab.nochim)/sum(ex_seqtab)
dim(ex_seqtab.nochim)
save(ex_seqtab.nochim, file = "./ex_seqtab.RData")
```

### 4 Balanced Dataset

```
balanced_dir <- "~/Downloads/" # CHANGE ME to location of file
balanced_fnF <- file.path(balanced_dir, "ERR777695_1.fastq.gz")
balanced_fnR <- file.path(balanced_dir, "ERR777695_2.fastq.gz")

ggsave(
  plotQualityProfile(balanced_fnF),
  filename = file.path(dirpath, "balanced_F_q.png") ,
  dpi = 300, width = 16, height = 10, units = "in")
ggsave(
  plotQualityProfile(balanced_fnR)  ,
  filename = file.path(dirpath, "balanced_R_q.png"),
  dpi = 300, width = 16, height = 10, units = "in")

balanced_filtF <- "balanced_F.fastq.gz"
balanced_filtR <- "balanced_R.fastq.gz"

balanced_out <- filterAndTrim(
  fwd = balanced_fnF, filt = balanced_filtF,
  rev = balanced_fnR, filt.rev = balanced_filtR,
  maxN=0, maxEE=2, truncQ=2,
  trimLeft=c(10,10), trimRight = c(40,40),
  compress=TRUE, verbose=TRUE)

balanced_errF <- learnErrors(balanced_filtF, multithread=TRUE)
balanced_errR <- learnErrors(balanced_filtR, multithread=TRUE)
balanced_dadaF <- dada(balanced_filtF, err=balanced_errF, multithread=TRUE)
balanced_dadaR <- dada(balanced_filtR, err=balanced_errR, multithread=TRUE)
balanced_mergers <- mergePairs(
  balanced_dadaF, balanced_filtF,
  balanced_dadaR, balanced_filtR, verbose=TRUE)

balanced_seqtab <- makeSequenceTable(balanced_mergers)

hist(nchar(colnames(balanced_seqtab)))
balanced_seqtab.nochim <- removeBimeraDenovo(
  balanced_seqtab, method="consensus",
  multithread=TRUE, verbose=TRUE)
sum(balanced_seqtab.nochim)/sum(balanced_seqtab)
dim(balanced_seqtab.nochim )
save(balanced_seqtab.nochim, file = "./balanced_seqtab.RData")
```

### 5 HMP

```
hmp_dir <- "~/Downloads/130403" # CHANGE ME to location of file
hmp_fnF <- file.path(hmp_dir, "Mock1_S1_L001_R1_001.fastq.bz2")
hmp_fnR <- file.path(hmp_dir, "Mock1_S1_L001_R2_001.fastq.bz2")

ggsave(
  plotQualityProfile(hmp_fnF),
  filename = file.path(dirpath, "hmp_F_q.png")  ,
  dpi = 300, width = 16, height = 10, units = "in")
ggsave(
  plotQualityProfile(hmp_fnR),
  filename = file.path(dirpath, "hmp_R_q.png")  ,
  dpi = 300, width = 16, height = 10, units = "in")
```

```r
# the reverse reads dont look great
hmp_filtF <- "hmp_F.fastq.gz"
hmp_filtR <- "hmp_R.fastq.gz"

hmp_out <- filterAndTrim(
  fwd = hmp_fnF, filt = hmp_filtF,
  rev = hmp_fnR, filt.rev = hmp_filtR,
  maxN=0, maxEE=2, truncQ=2,
  truncLen=c(240,200), trimLeft=c(20,20),
  compress=TRUE, verbose=TRUE)

hmp_errF <- learnErrors(hmp_filtF, multithread=TRUE)
hmp_errR <- learnErrors(hmp_filtR, multithread=TRUE)
hmp_dadaF <- dada(hmp_filtF, err=hmp_errF, multithread=TRUE)
hmp_dadaR <- dada(hmp_filtR, err=hmp_errR, multithread=TRUE)
hmp_mergers <- mergePairs(
  hmp_dadaF, hmp_filtF,
  hmp_dadaR, hmp_filtR, verbose=TRUE)

hmp_seqtab <- makeSequenceTable(hmp_mergers)

hist(nchar(colnames(hmp_seqtab)))
hmp_seqtab.nochim <- removeBimeraDenovo(
  hmp_seqtab, method="consensus",
  multithread=TRUE, verbose=TRUE)
sum(hmp_seqtab.nochim)/sum(hmp_seqtab)
dim(hmp_seqtab.nochim )
save(hmp_seqtab.nochim, file = "./hmp_seqtab.RData")
```

### 6 Combining sequence tables

Having processed all the datasets without DADA2, the tables were merged.

```r
if (SPEED){
  for (p  in c("hmp_seqtab.RData", "ex_seqtab.RData",
               "balanced_seqtab.RData", "endo_seqtab.RData")){
    load(p)
  }
}

# Now, we format them so we can bind them all together
#View(hmp_seqtab.nochim)
#View(endo_seqtab.nochim)

rownames(hmp_seqtab.nochim) <- "HMP"
rownames(ex_seqtab.nochim) <- "Extremes"
rownames(balanced_seqtab.nochim) <- "Balanced"

thesenames <- c(
  rownames(endo_seqtab.nochim),
  rownames(hmp_seqtab.nochim),
  rownames(ex_seqtab.nochim),
  rownames(balanced_seqtab.nochim)
)
combined_seqtab <- dplyr::bind_rows(
  as.data.frame(endo_seqtab.nochim, row.names = rownames(endo_seqtab.nochim)),
  as.data.frame(hmp_seqtab.nochim,  row.names = rownames(hmp_seqtab.nochim)),
  as.data.frame(ex_seqtab.nochim,  row.names = rownames(ex_seqtab.nochim)),
```

```
  as.data.frame(balanced_seqtab.nochim,  row.names = rownames(balanced_seqtab.nochim)),

)
rownames(combined_seqtab) <- thesenames
combined_seqtab <- as.matrix(combined_seqtab)
#combined_seqtab <-  combined_seqtab[, 1:40]
```

### 7 Recreating the DADA2-formatted SILVA and augmented databases

Now, we need to reformat the SILVA database for use with DADA2. We could use the pre-built one, but as we have to build two (the normal SILVA 132 and our augmented one), we outline the steps here.

The taxonomy and reference alignment files are downloaded:

```
wget https://www.arb-silva.de/fileadmin/silva_databases/release_132/Exports/SILVA_132_SSURef_tax_silva.fasta.gz
wget https://www.arb-silva.de/fileadmin/silva_databases/release_132/Exports/SILVA_132_SSURef_Nr99_tax_silva_full_al
```

We recreate the databases described here: https://zenodo.org/record/1172783, but to do that, we need to add our sequences to the alignment, which we can do with MAFFT.

```
mkdir ~/Downloads/dada2_dbs/

zcat < ~/Downloads/SILVA_132_SSURef_tax_silva.fasta.gz | \
  sed 's/>/> /g' | cut -f 1,3 -d " " | sed 's/ //g'  | \
  awk '/^>/ {$0=$0 ";"}1' |gzip -c > ~/Downloads/dada2_dbs/silva.132.formated.fasta.gz
zcat < ~/Downloads/dada2_dbs/silva.132.formated.fasta.gz | \
  awk 'BEGIN{RS=">";FS="\n"}NR>1{printf ">%s\n",$1;for (i=2;i<=NF;i++) {gsub(/U/,"T",$i); printf "%s\n",$i}}' | \
  gzip -c > ~/Downloads/dada2_dbs/silva.132.dna.formated.fasta.gz

cp  ~/Downloads/SILVA_132_SSURef_tax_silva.fasta.gz  ~/Downloads/SILVA_132_SSURef_tax_silva_plus.fasta.gz
cat ./results/fast_focusDB_ribo16s.fasta |gzip -c  >>  ~/Downloads/SILVA_132_SSURef_tax_silva_plus.fasta.gz

zcat < ~/Downloads/SILVA_132_SSURef_tax_silva_plus.fasta.gz | \
  sed 's/>/> /g' | cut -f 1,3 -d " " | sed 's/ //g'  | awk '/^>/ {$0=$0 ";"}1'  | \
  gzip -c > ~/Downloads/dada2_dbs/silva.132.plus.formated.fasta.gz
zcat < ~/Downloads/dada2_dbs/silva.132.plus.formated.fasta.gz | \
  awk 'BEGIN{RS=">";FS="\n"}NR>1{printf ">%s\n",$1;for (i=2;i<=NF;i++) {gsub(/U/,"T",$i); printf "%s\n",$i}}' |\
  gzip -c > ~/Downloads/dada2_dbs/silva.132.plus.formated.dna.fasta.gz
```

Now, we can use DADA2's built-in command to create a species level databases for both:

```
dada2:::makeSpeciesFasta_Silva(
  "~/Downloads/SILVA_132_SSURef_tax_silva.fasta.gz",
  "~/Downloads/dada2_dbs/silva_species_assignment_v132.fa.gz")
dada2:::makeSpeciesFasta_Silva(
  "~/Downloads/SILVA_132_SSURef_tax_silva_plus.fasta.gz",
  "~/Downloads/dada2_dbs/silva_plus_species_assignment_v132.fa.gz")
#313502 sequences with genus/species binomial annotation output.
#319084 sequences with genus/species binomial annotation output.
```

### 8 Assigning taxonomy to the combined sequence table

Next, we assign taxa to the merged ASV table. Matrices for the data with both the SILVA db and our SILVA+ db created with focusDB. This is then used to determine whether the taxonomic assignment changes between the two databases.

```
taxa_silva  <- assignTaxonomy(
  combined_seqtab,
  "~/Downloads/dada2_dbs/silva.132.dna.formated.fasta.gz",
  multithread=TRUE, verbose = T)

taxa_silva_species <- assignSpecies(taxa_silva,verbose = T, "~/Downloads/dada2_dbs/silva_species_assignment_v132.fa
# 710 out of 4098 were assigned to the species level.

taxa_silva_plus  <- assignTaxonomy(
  combined_seqtab,
  "~/Downloads/dada2_dbs/silva.132.plus.formated.dna.fasta.gz",
  multithread=TRUE, verbose = T)

taxa_silva_plus_species <- assignSpecies(
  taxa_silva_plus,
  "~/Downloads/dada2_dbs/silva_plus_species_assignment_v132.fa.gz",
  allowMultiple = T, verbose = T)
# 713 out of 4098 were assigned to the species level.

save.image("taxa_assigned.RData")
```

```
if (SPEED){load("taxa_assigned.RData")}
together <- merge(
  by="seq", all=TRUE,
  data.frame(silva=paste(
    as.character(taxa_silva_species[,1]),
    as.character(taxa_silva_species[,2])),
    seq = row.names(taxa_silva_species),
    stringsAsFactors=FALSE),
  data.frame(focuDB=paste(
    as.character(taxa_silva_plus_species[,1]),
    species=as.character(taxa_silva_plus_species[,2])),
    seq = row.names(taxa_silva_plus_species),
    stringsAsFactors=FALSE)
)
#together[together$silva!=together$focuDB, ] %>% View()
write.table(together[together$silva!=together$focuDB, ],
            file = "STABLE_different_assignment.tab", row.names = F, sep="\t")
```
